# Maternal total sleep deprivation causes oxidative stress and mitochondrial dysfunction in oocytes associated with fertility decline in mice

**DOI:** 10.1101/2024.06.13.598954

**Authors:** Zi-Yun Yi, Qiu-Xia Liang, Qian Zhou, Lin Yang, Qing-Ren Meng, Jian Li, Yi-hua Lin, Yan-pei Cao, Chun-Hui Zhang, Heide Schatten, Jie Qiao, Qing-Yuan Sun

## Abstract

Previous studies have shown sleep deprivation has an impact on several aspects of health and disease. Sleep deprivation suppresses melatonin production and excessive HPA activation in women, which may reduce the chances of fertility. However, little is known about how and by what relevant mechanisms it affects female fertility. In this study, female mice underwent 72 hours of total sleep deprivation (TSD) caused by rotating wheel or 2 different controls: a stationary wheel, or forced movement at night. Even though, there was no evident anomaly of natural ovulation, the TSD female mice showed impaired early embryo development . Overall levels of estrogen and FSH were lower throughout the estrus cycle (especially in proestrus). RNA Sequencing (RNA-Seq) was used to discover differentially expressed genes (DEGs) that might be involved in oocyte quality after TSD. A total of 42 genes showed significant differential expression in GV oocytes after TSD. These included genes such as *Usmg5, Atp5k, Ccnd2* and *Tpm3*, which were enriched in gene ontology terms of mitochondrial protein complex, oxidoreductase activity, cell division, cell cycle G1/S phase transition, as well as others. The increased concentrations of reactive oxygen species (ROS) in germinal vesicle (GV) and metaphase II (MII) oocytes from TSD mice were observed, which might be induced by impaired mitochondrial function caused by TSD. The GV oocytes displayed increased mitochondrial DNA (mtDNA) copy number and a significant transient increase in inner mitochondrial membrane potential (△ψm) from the TSD mice probably due to compensatory effect. In contrast, MII oocytes in the TSD group showed a decrease in the mtDNA copy number and a lower △ ψm compared with the controls. Furthermore, abnormal distribution of mitochondria in the GV and MII oocytes was also observed in TSD mice, suggesting mitochondrial dysfunction. In addition, abnormal spindle and abnormal arrangement of chromosomes in MII oocytes were markedly increased in the TSD mice compared with the control mice. In conclusion, our results suggest that TSD significantly alters the oocyte transcriptome, contributing to oxidative stress and disrupted mitochondrial function, which then resulted in oocyte defects and impaired early embryo development in female mice.

## Introduction

Sleep is a biological phenomenon observed in most species, and It is necessary for proper physical and mental performance [1]. Sleep loss is a condition that is prevalent in people who have a hectic life style and 20 percent of the adult population is affected by Sleep deficiency [2]. Sleep deprivation impacts negatively on the nervous system [3], immune system [4], endocrine system, cardiovascular system [5,6], as well as the male reproductive system [7]. Acute sleep deprivation leads to endocrine changes, with decreased concentrations of testosterone (T) and E2, adversely affecting male sexual function and sperm concentration, quantity and quality [7]. In fact, sleep disturbances such as insomnia are common complaints among women [8]. Hormonal fluctuations in the menstrual cycle have been linked to sleep deprivation in female rats [8].

Short sleep duration is also associated with irregular menstrual cycles, which may harm reproductive health in human [9]. Although many studies have shown that Sleep deprivation affects several aspects of health and diseases, little is known about how the female reproductive system is affected by this condition. Mitochondria are the main energy generators in oocyte ooplasm, and they crucially affect oocyte quality and early embryonic development [8,10]. Mitochondrial complement size, mitochondrial DNA (mtDNA) copy number, and spatial distribution have been considered keys to the development ability of oocyte and embryo development, including normal spindle organization and chromosomal segregation, timing of cell cycle progression, and morpho-dynamic processes such as compaction, cavitation and blastocyst hatching [11,12]. Thus, mitochondrial dysfunctions, resulting in reduced oxidative phosphorylation (OXPHOS) and decreased ATP production, decreased oxidative capacity and antioxidant defense, effects which could in turn be involved in reduced the quality and developmental potential of oocytes and early embryos [13,14]. Recently, it was reported that one night of sleep curtailment in humans acutely induced inefficient mitochondrial function and increased the levels of acylcarnitines in plasma, which are necessary intermediates for mitochondrial fatty acid oxidation (FAQ) [15]. In addition, in sleep-deprived mice, oxidative stress and acetylation of mitochondrial proteins are increased in parallel with mitochondrial injury, apoptosis is activated, and Locus ceruleus neurons (LCns) are lost [3]. More importantly, paradoxical sleep deprivation can decrease the activity of complex I-II and III enzymes of the mitochondrial electron transport chain, which may further lead to increased mitochondrial dysfunction and decreased availability of brain energy [16].

ROS, superoxide and H_2_O_2_ are products of mitochondria during the electron transfer process that occurs along the electron transport chain (ETC) [6]. Once Dysfunctional mitochondria produce ROS levels that exceed antioxidant defenses, redox homeostasis is damaged and oxidative stress is induced, which may eventually induce dysfunction and apoptosis of oocytes and embryos [17]. Sleep deprivation has also been proved to relate to elevated levels of reactive oxygen species (ROS) [18]. Sleep deprivation involves elevated energy expenditure to maintain electrical potentials, which requires a significant amount of oxygen, resulting in a large number of ROS [18]. In addition, sleep deprivation leads to oxidative damage, promotes mitochondrial dysfunction and ROS production, exacerbating the oxidative stress (OS) states in many species [6,18]. Melatonin (N-acetyl5-methoxytryptamine) is a major pineal secretory product that exists at a high level in human ovarian follicular fluid [19]. A low level of follicular melatonin induced the accumulation of excessive ROS in oocytes, leading to meiotic failure and the production of aneuploid eggs in infertile women such as aged women [20], and obesity women [21]. Sleep deprivation alters the secretion of endogenous melatonin, which may expose the follicles to high levels of oxidative stress [19]. Therefore, in this study, we assume that sleep deprivation induces mitochondrial dysfunction and oxidative stress in oocytes, which could in turn result in oocyte dysfunctions and impaired early embryo development.

Sleep disorders also affect many aspects of physiological processes, by regulating the expression of specific genes and synthesis and secretion the of related proteins [1]. Sleep deprivation could decrease sperm motility and cause male infertility by regulating the expression of Interleukin (IL6), nitric oxide synthase (INOS), and tumor necrosis factor alpha (TNFα) [1,22]. Oocyte transcription is silenced after germinal vesicle rupture (GVBD), so the final stages of oocyte meiotic maturation and early embryo development depend on protein synthesis from stored transcripts [23]. Here we also wondered if sleep deprivation could cause alteration of gene expression in oocytes.

To address the above problems, we used a rotating drum method-induced total sleep deprivation (TSD) mouse model to investigate the effects of total sleep deprivation on female mouse reproduction. We also employed single GV oocyte RNA sequencing (RNA-Seq) to detect changes in transcript abundance after sleep deprivation. We found that the levels of two important sex hormones (estrogen and FSH) were affected. Further study identified that TSD did affect oocyte quality and early embryo development by altering gene expression in the oocytes, causing oxidative stress and mitochondrial dysfunction, inducing spindle and chromosome defects in oocytes.

## Results

### The body weight is normal, but the sex hormone levels are disturbed in TSD mice

To explore the effect of maternal total sleep deprivation (TSD) on female mouse fertility, the mouse TSD model was induced in mice by the rotating drum method [5,24]. As shown in Figure 1 A-B, the body weights at the beginning (day 0) and body weight changes from the beginning (day 0) to termination (day 3 of total sleep deprivation experiment) of the sleep deprivation experiment were similar in all groups. We measured the E2 and FSH levels at different stages of the estrus cycle in the stationary, TSD and forced night activity groups. We found that the E2 level decreased at all stages of the estrus cycle in the TSD mice compared with that of the stationary or the forced movement groups (Fig. 1C). The FSH level was significantly decreased at proestrus in the TSD group, while it was similar at other stages of the estrus cycle in all groups (Fig. 1D).

**Figure1.**
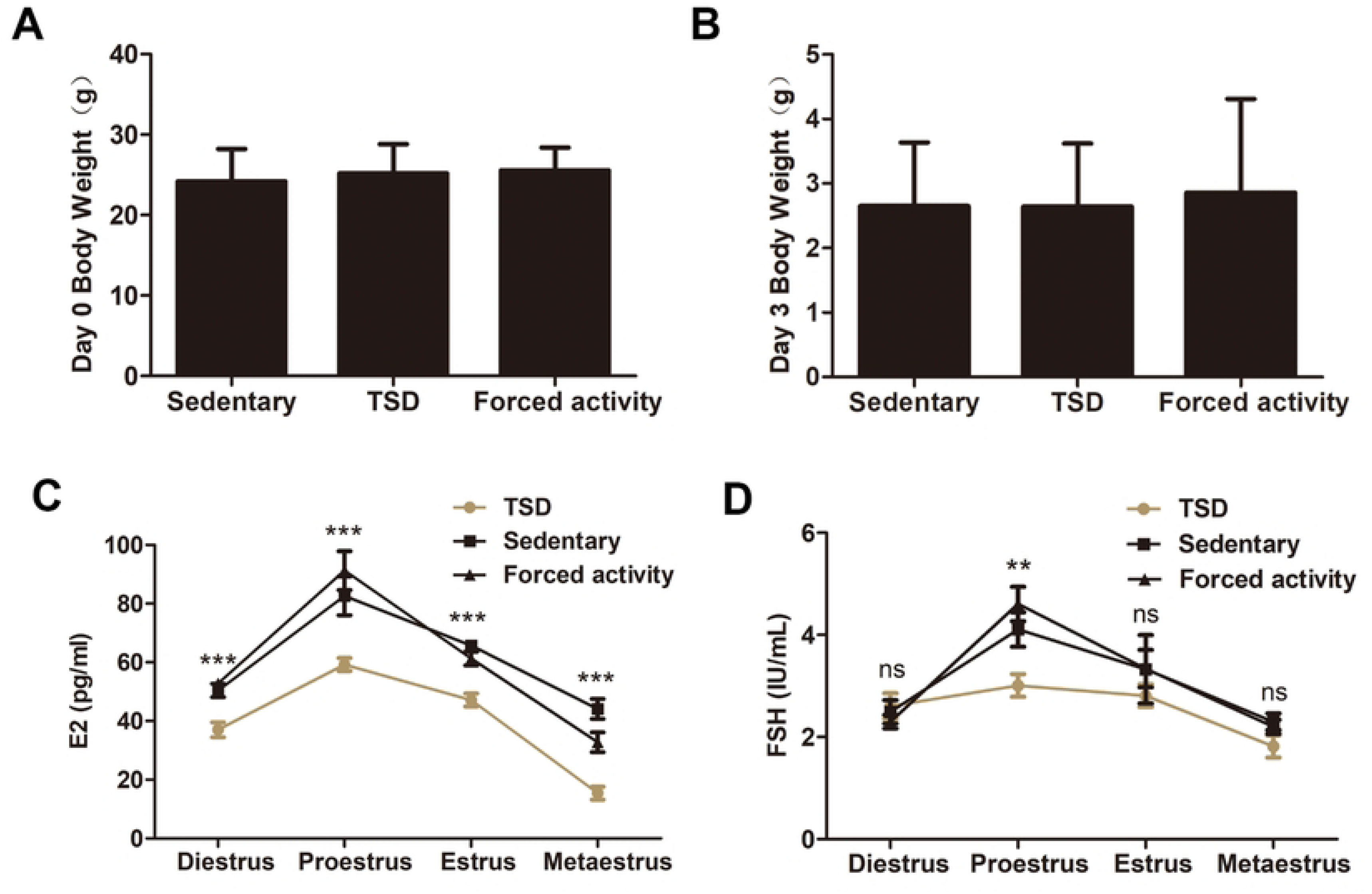
The effects of sleep deprivation on maternal weight and sex hormone levels. (**A** and **B**) Body weights at the beginning and body weight changes from beginning to termination of the sleep deprivation experiment in the stationary group, the TSD group and the forced movement group. The data are expressed as means ± SEM. **(C** and **D**) Concentrations of serum estrogen (C) and FSH (D) during four stages of estrus cycle in the stationary, forced night activity, and TSD groups. The data are expressed as means ± SEM. ***, P<0.001; **, P<0.01; NS, no significant difference.

### The TSD mice display subfertility, but normal number of ovulated eggs

As shown in Figure 2A, the natural ovulation test showed no significant difference in ovulation in TSD mice compared with the stationary and the forced night activity control groups. The TSD mice ovulated at an average of 17. 5±1.24 eggs, whereas the stationary and forced movement mice ovulated at an average of 18.91±0.78 and 19.3±0.86 eggs, respectively (Fig. 2A). The fertilization rate of natural ovulated eggs showed no significant difference compared with the stationary and the forced night activity control groups (Figure 2B).

**Figure 2.**
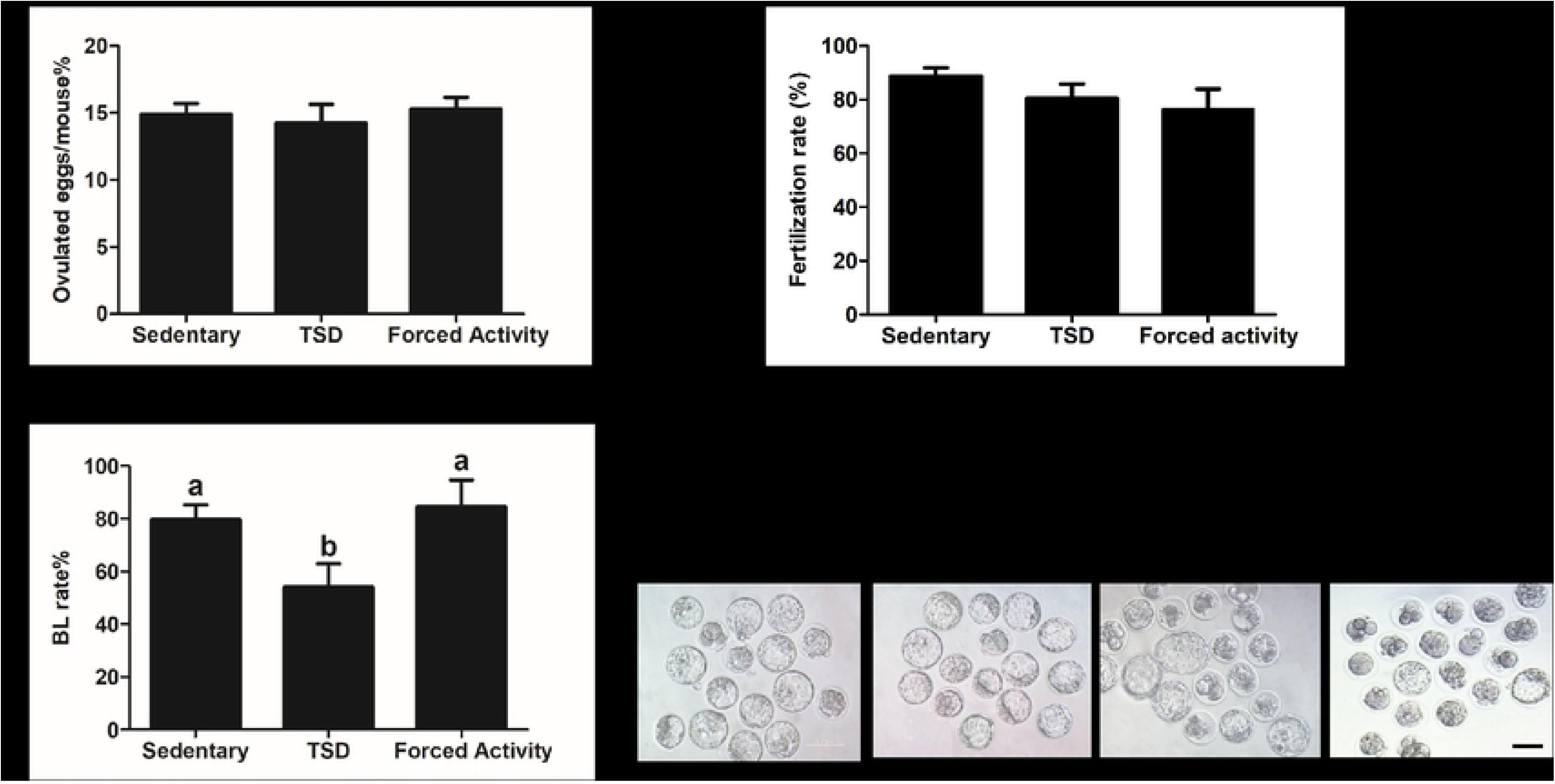
The total sleep deprivation (TSD) mice ovulated normal number of eggs but showed impared early embryo development . **(A**) The average number of ovulated eggs. Four female mice with plugs were examined for each group. **(B)** The fertilization rate for the stationary group, the TSD group and the forced movement group. The data are expressed as means± SEM of at least three replicates. Different letters indicate statistically significant difference (P<0.05). (**C**) The blastocyst rates of fertilized eggs after in vitro culture (IVC) for 5d. (**D**) Development of embryos after 5 days of culture in the stationary, the TSD and the forced movement groups. Bar=100μm. In A and C, data are expressed as means ± SEM of three replicates. Different letters indicate statistically significant difference (P<0.05).

### Total sleep deprivation affects early embryo development

We further defective determined preimplantation embryo development in vitro. Fertilized eggs were collected from the oviduct of female mice with plugs. The blastocyst rates in the TSD group was significantly decreased compared to the control groups (54±8.96% versus 79.61±5.597% and 84.55±10.12%, control; P<0.05) (Fig. 2 C-D) (P<0.05). Taken together, total sleep deprivation affects early embryonic development.

### Total sleep deprivation alters gene expression of oocytes

To understand the molecular mechanism underlying impaired embryonic development after sleep deprivation, we performed RNA-Seq to analyze the transcription profiles of mouse oocytes from the TSD group and two control groups (Fig.3). A total of 64 DEGs were differentially expressed between stationary and TSD oocytes (with more than 2-fold changes, p<0.05) (Fig. 3A; SETa). While there were 109 DEGs obtained between the stationary and forced night activity groups (SETb). And there were 42 DEGs obtained by subtracting SETb from SETa, namely eliminating the effects of gentle exercise on the transcriptome, and the 42 DEGs were only caused by TSD. Subsequent analysis was based on the 42 DEGs (DEG_TSD_). Table 1 shows the 12 significantly regulated genes. We found that the expressions of genes previously reported to be related to mitochondrial function, ATP synthase, cell cycle G1/S phase translation, and cell division, such as *Usmg5* [25], *Atpk5* [26], *Ccnd2* [27] and *Tpm3* [28], were altered. To validate the RNA-Seq data, the RT-PCR was performed on GV oocyte samples from the stationary, TSD, and forced night activity groups, respectively (Fig. 3, C-D). Although the folding changes of RT-PCR and RNA-Seq were different, the expression trends of the selected genes were similar.

**Figure 3.**
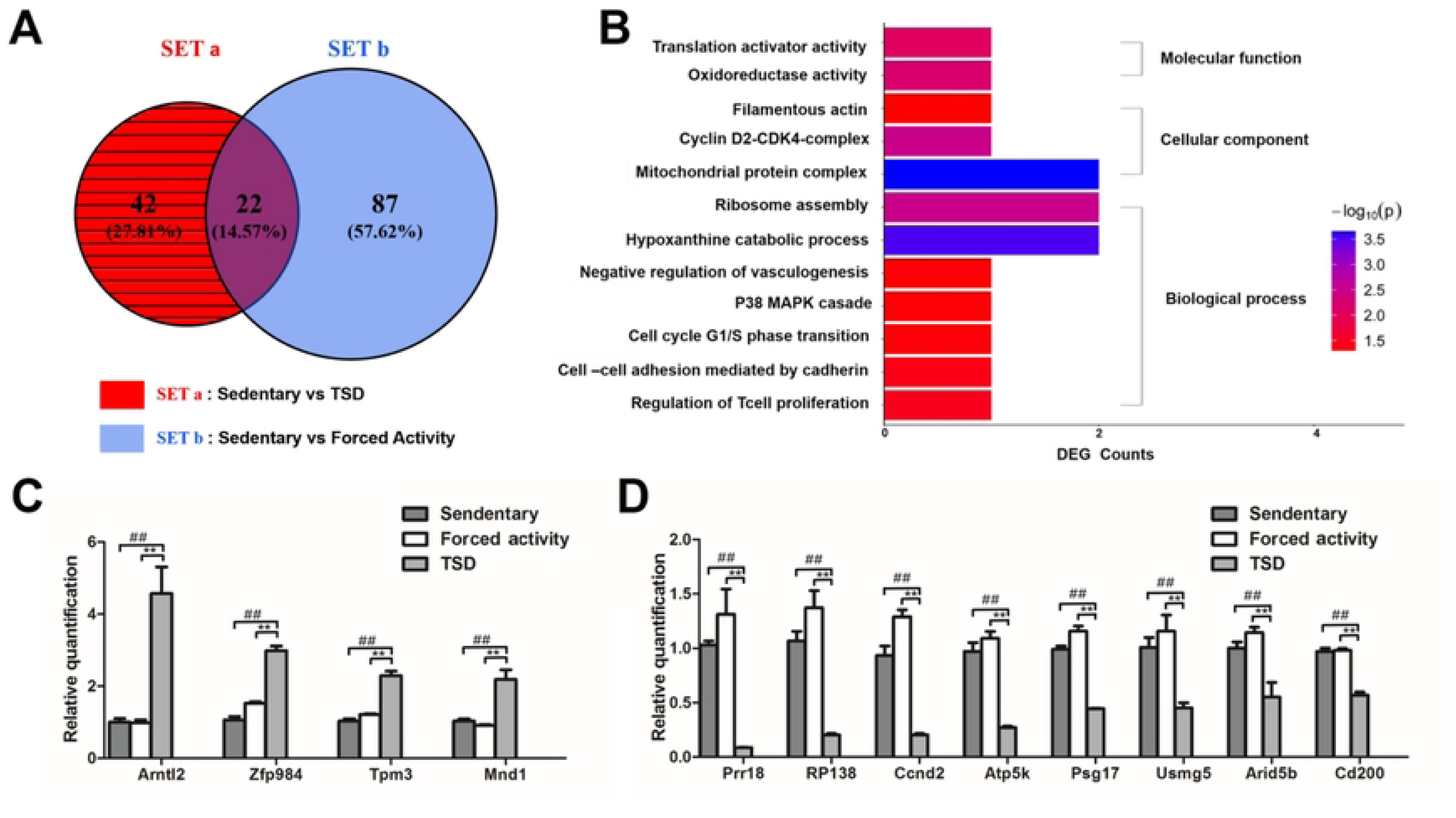
Single-GV oocyte RNA sequencing. (**A**) The Venn diagram of DEGs in stationary and forced night activity groups separately compared with those in the TSD group. SETa is the 64 DEGs between the TSD and stationary groups, SETb is the 109 DEGs between the stationary and forced night activity groups, and DEG_TSD_ defines the 42 DEGs (the shaded part) obtained by subtracting SETb from SETa, namely the DEGs caused by Total Sleep Deprivation (DEG_TSD_) alone and not by exercise. Subsequent analysis was based on DEG_TSD._ **(B)** Selected GO analysis of 42 DEGs found by RNA-Seq including biological process, molecular functions and cellular components. The horizontal axis shows the count of genes, and the color of bar represents –Log10 (p-value). Description of each GO term was marked in the bar plot. Their p-values were all less than 0.05. **(C and D)** RT-PCR to validate selected 12 differentially expressed genes. **(C)** The relative quantification of the 4 up-regulated genes. **(D)** The relative quantification of the 8 down-regulated genes. Gray bars: the stationary control group; white bars: the forced night activity control (FA); light grey bars: the total sleep deprivation (TSD) group. The data were expressed as mean± SEM of at least three replicates. ##P<0.05: significant difference between the stationary group and the TSD group; ** P <0.05: significant difference between the forced night activity group and the TSD group.

Gene ontology (Go) enrichment analysis was used to study the effects of TSD on biological functions. Go analysis on 35 annotated DEGs of the 42 total DEG_TSD_ identified a number of important functions (Fig. 3, B), such as mitochondrial protein complex, oxidoreductase activity, p38 MAPK cascade, filamentous actin and translation activity, cell cycle G1/S transition, as well as others.

### The intra-cellular ROS level of the GV and MII oocytes of the TSD mice increased

To estimate intracellular oxidative stress in oocytes, we measured the level of ROS levels in the GV and MII oocytes (Fig. 4A). The DCFA-DA fluorescence intensity of each group was evaluated (Fig. 4, B-C). The DCFA-DA fluorescence intensities were significantly increased in the GV and MII oocytes of the TSD group compared to those in the stationary and the forced night activity groups, indicating a higher level of ROS. In addition, as shown in Figure 4A, ROS in TSD mouse MII oocytes are distributed in clumps in the cytoplasm. Therefore, total sleep deprivation induces an increase in oxidative stress in oocytes.

**Figure 4.**
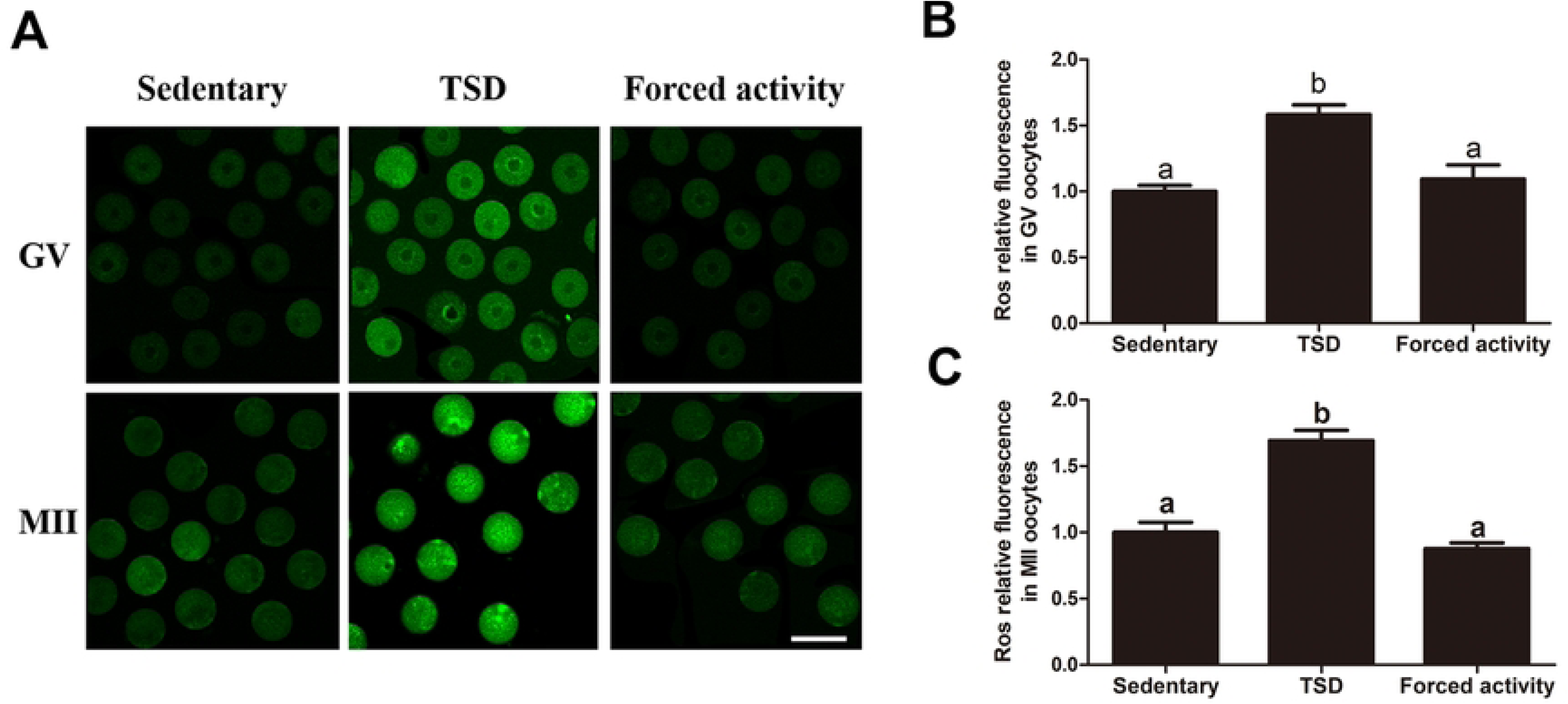
Effects of total sleep deprivation on ROS in the GV and MII oocytes. **(A)** Representative images of carboxy-H2DCF fluorescence in the GV and MII oocytes. The fluorescence intensity was significantly higher in the GV and MII oocytes of the TSD group compared to that of the stationary and the forced night activity groups. Bar=100μm. **(B)** The fluorescence intensity of the carboxy-H2DCF fluorescence of each GV oocyte in the stationary, the TSD and the forced night activity groups were quantified using ZEN (2012) software. Data are expressed as mean ± SEM of at least 3 independent experiments and 3 mice were killed to obtain a minimum of 50 oocytes for each experiment. Different letters indicated statistically significant differences (P<0.05). **(C)** Effects of total sleep deprivation on ROS production measured by the carboxy-H2DCF fluorescence of each of the MII oocytes using ZEN (2012) software. Data are mean ± SEM of at least 3 independent experiments. Four superovulated mice were killed to obtain a minimum of 30 oocytes for each experiment. Different letters indicate statistically significant differences (P<0.05).

### Increased mtDNA copy number in the GV oocytes but decreased mtDNA in the MII oocytes in the TSD mice

Because the functional state of mitochondria affects the quality of oocytes and contributes to the process of fertilization and embryonic development, mitochondria are directly involved in the reproductive process at several levels. Mitochondria play a major role in cellular energetic metabolism, homeostasis, and death.

They have their own multi-copy genome, which is inherited maternally. Mitochondrial DNA (mtDNA) content directly influenced mitochondrial function during oogenesis and early embryonic development [29]. Here, we investigated whether maternal sleep deprivation affected mtDNA content in the GV and MII oocytes. Quantitative RT-PCR was performed on single fully-grown GV and MII oocytes in all groups. As shown in Figure 5A, GV oocytes in the TSD group contained more mtDNA copies compared to those in the stationary and the forced night activity groups (474600±23000 versus 317300±29240 and 362800±32150, control; P<0.05) probably due to compensatory effect for their survival and further maturation. However, the average mtDNA copy number in MII oocytes from the TSD mice was significantly decreased compared with that of the MII oocytes from the stationary control and the forced night activity control groups (303300±22680 versus 390000±25830 and 427300±43440, control; P<0.05) (Fig. 5B). This phenomenon was also observed in our previous report on mtDNA copy number changes in oocytes derived from insulin-resistant mouse model [13].

**Figure 5.**
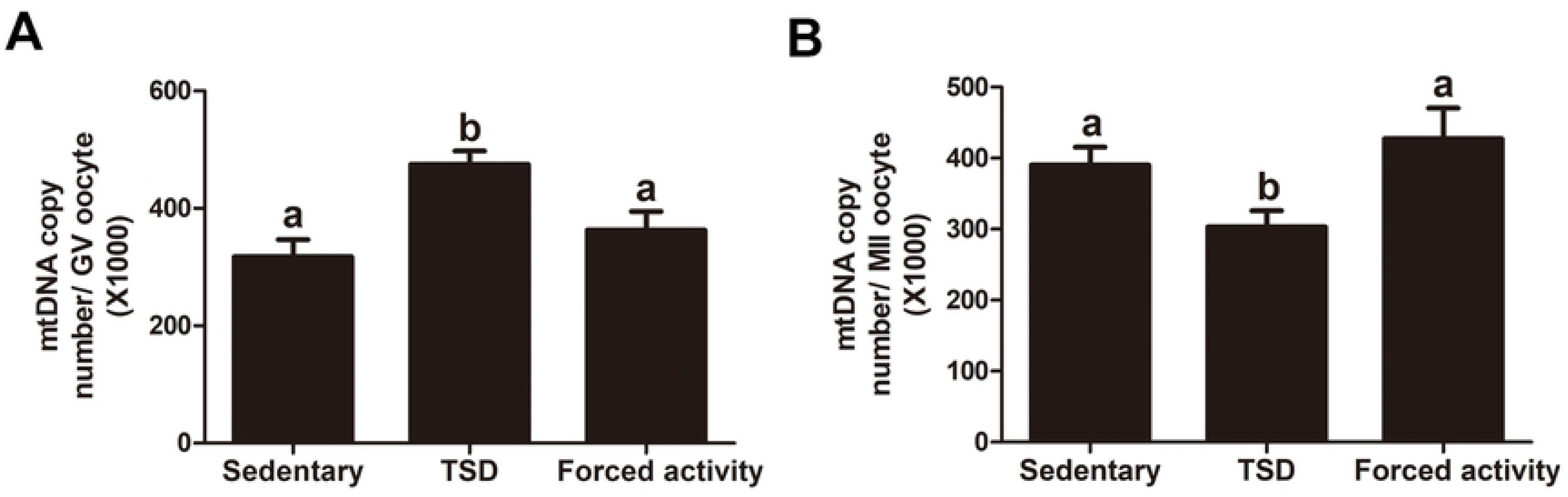
Effects of sleep deprivation on the mtDNA copy number in single GV and MII oocytes. Single oocyte was extracted and used for determining the mtDNA copy number. (A) The mtDNA copy number in single GV oocyte from the stationary, TSD and forced night activity groups. (B) The mtDNA copy number in single MII oocyte from the stationary, TSH and forced night activity groups. Data are expressed as mean ± SEM and at least 30 oocytes were examined for each group. Different letters indicate statistically significant differences (P<0.05).

### Maternal total sleep deprivation disrupts mitochondrial redistribution in the oocytes

Studies in some mammalian species have shown that the distribution of mitochondria undergoes stage-specific changes during oocyte maturation and early embryogenesis [10,13]. It is thought that spatial remodeling of mitochondria may lead to elevated environmental ATP levels in cytoplasmic regions where stage-specific activities may require higher energy [10,11]. We next examined whether maternal sleep deprivation influences the distribution of mitochondria by fluorescence staining. The data showed that the majority of the GV oocytes in the stationary and the forced night activity groups (70±1% and 65±5%) showed a perinuclear mitochondrial distribution while the portion of perinuclear distribution decreased in the TSD group (42±2%,P<0.05). Whereas, the proportions of homogeneous distribution pattern were similar between the stationary, the TSD and the forced movement groups (25±2%, 30±5% and 29±3%, P>0.05). Importantly, oocytes from the TSD mice displayed a much higher proportion of aggregating or clustering mitochondrial distribution compared to the stationary and the forced night activity groups (28±6% versus 5±1% and 6±2 %, control; P<0.05) (Fig. 6, A-B).

**Figure 6.**
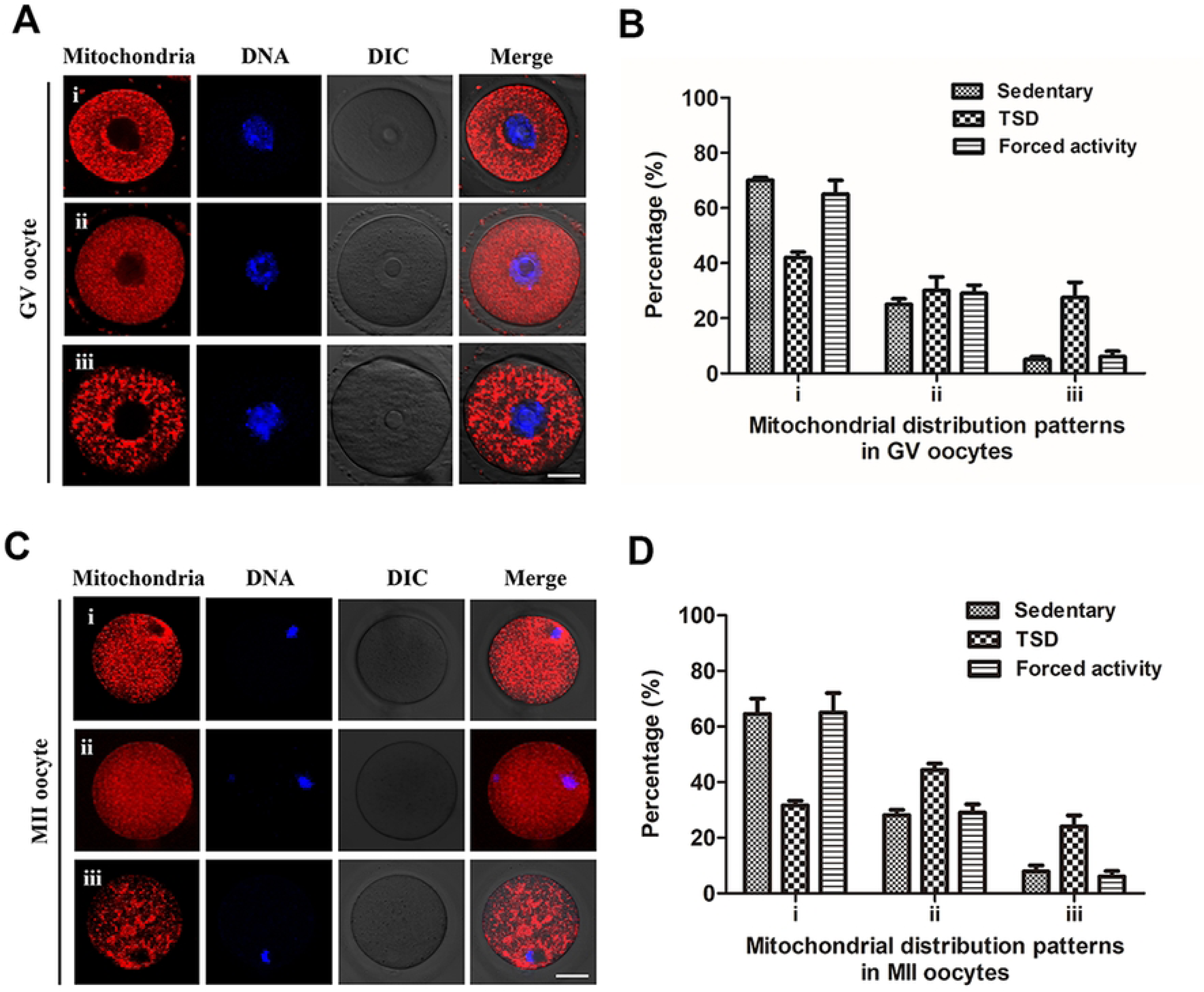
Maternal sleep deprivation disrupts mitochondrial redistribution in the GV and MII oocytes. The oocytes were stained with MitoTracker-Red to detect the mitochondrial distribution patterns, and chromosomes were counterstained with Hoechst 33342 (blue) to confirm meiotic stages. **(A)** In the GV oocytes, three different distributions of mitochondria were observed: (i) perinuclear distribution, (ii) homogeneous distribution and (iii) clustered distribution. Bar=20μm. **(B)** Quantification of the GV oocytes with each mitochondrial distribution pattern from the stationary, TSD and forced movement groups. **(C)** In ovulated MII oocytes, three mitochondrial distribution patterns were detected: (i) perinuclear distribution, (ii) homogeneous distribution and (iii) clustered distribution. Bar=20μm. **(D)** Proportions of MII oocytes from stationary, TSD and forced night activity groups to show each mitochondria distribution pattern described in C. Data in B and D are expressed as mean ± SEM of at least 3 independent experiments. Three mice were killed to obtain a minimum of 50 oocytes to investigate the mitochondria distribution in the GV oocyte for each replicate experiment, and 4 superovulated mice were killed to obtain a minimum of 30 oocytes to investigate the mitochondrial distribution in the MII oocytes for each independent replicate.

Compared with the two controls, the distribution pattern of polarized mitochondria of MII oocytes in TSD group was reduced (32±2% versus 64±5% and 65±7%, control; P<0.05)(Fig. 6, C-D), while the distribution of smooth and homogeneous mitochondria increased in the TSD group compared with the two control groups (44±2% versus 28±2% and 29±3%, control; P<0.05). As shown in Figure 7 D, the proportion of the granular and clumped mitochondrial distribution also increased in the TSD group relative to the two control groups (24±4% versus 8±2% and 6±2%, control; P<0.05). These results suggest that total sleep deprivation leads to insufficient spatial remodeling of mitochondria during oocyte maturation.

**Figure 7.**
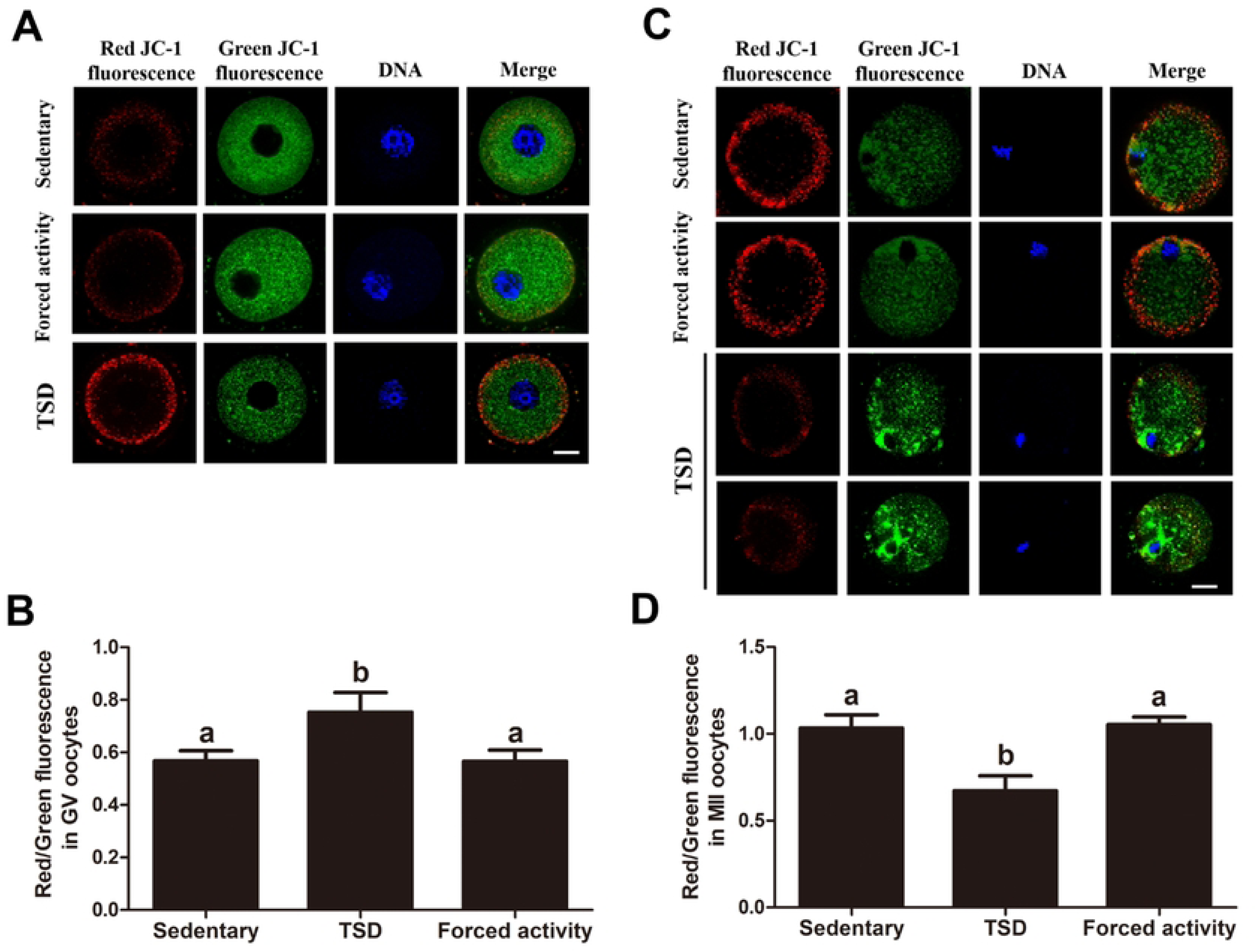
Effects of sleep deprivation on the mitochondrial transmembrane potential in the GV and MII oocytes. (A) Images of the GV oocytes in stationary, TSD and forced night activity groups stained for JC-1. JC-1 is a fluorescent dye that emits a red fluorescence as JC-1 aggregates in high-polarized mitochondria and emits a green fluorescence as JC-1 monomer in low-polarized mitochondria. Bar=20μm. (B) The ratio of red/green fluorescence intensities in the GV oocytes from the stationary, TSD and forced night activity groups were quantified as an indicator of mitochondrial activity. The data represents means ± SEM of at least three independent experiments. Three mice were killed to obtain a minimum of 50 GV oocytes for each replicate experiment. Different letters denote statistically significant differences (P<0.05). (C) Images of MII oocytes in stationary, TSD and forced night activity groups stained by JC-1 probe. Bar=20μm. (D) The ratio of red/green fluorescence intensity in ovulated MII oocytes was examined in all groups. The data represents means ± SEM of at least three independent experiments. Four superovulated mice were killed to obtain a minimum of 30 MII oocytes for each replicate experiment. Bars with different letters differ significantly (P<0.05).

### Maternal total sleep deprivation affects the inner mitochondrial membrane potential in oocytes

At an inner mitochondrial membrane potential (△ψm) <100mV, JC-1 accumulates as multimers that emit a green fluorescence in the FITC channel (low-polarized mitochondria). At △ψm>140 mV, JC-1 forms J-aggregates and gives off a red fluorescence in the rhodamine isothiocyanate channel [30]. Thus, an increase in the red/green fluorescence intensity ratio indicates an increase in mitochondrial activity [31]. As shown in Figure 8, we investigated whether maternal total sleep deprivation altered mitochondrial activity in oocytes by examining the red: green fluorescence ratio [32].

**Figure 8.**
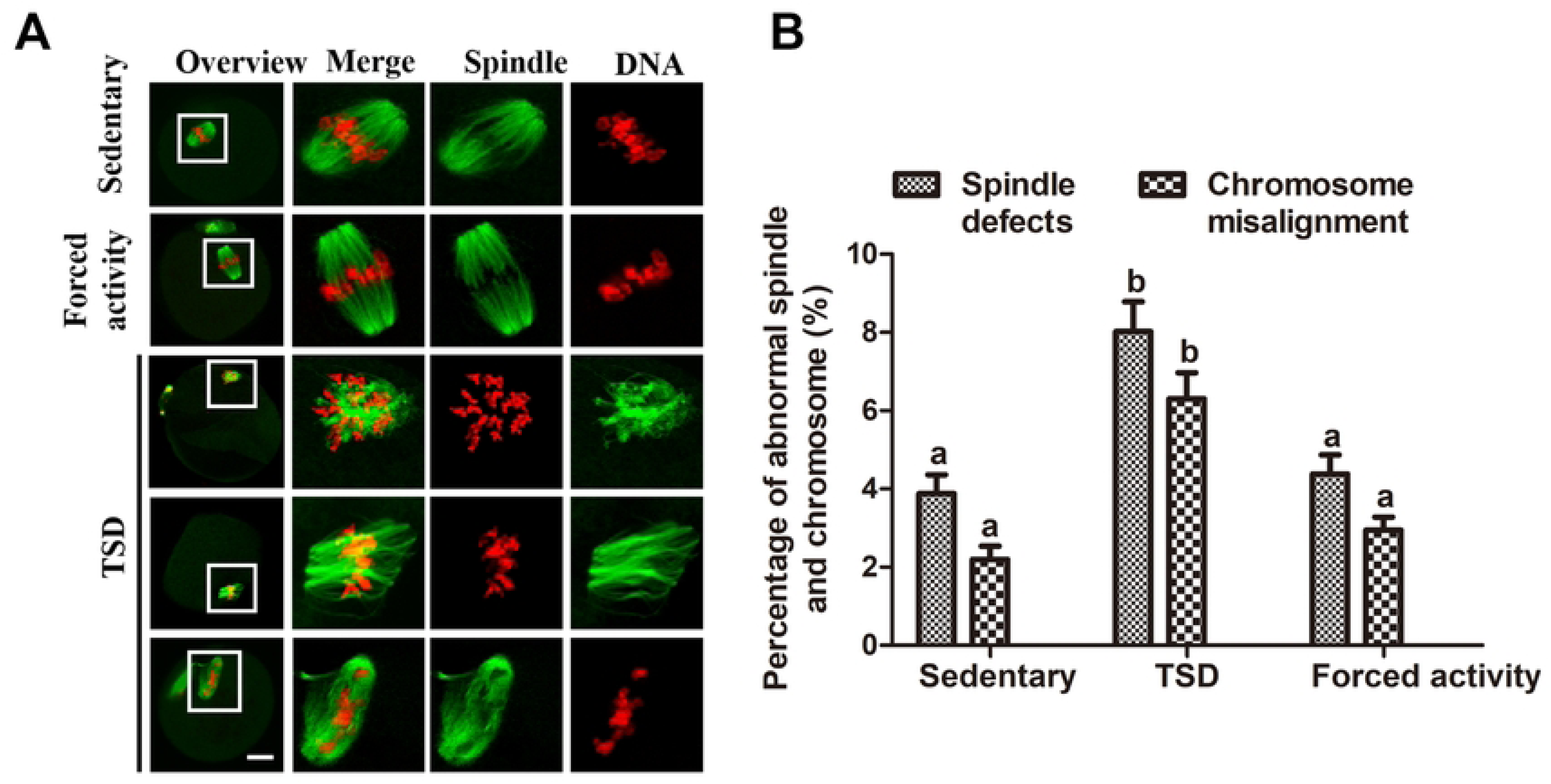
Maternal sleep deprivation leads to defective spindles and misaligned chromosomes. **(A)** Representative images of the spindle and chromosomes in stationary, TSD and forced night activity groups. MII oocytes from the stationary and the forced movement mice present a normal spindle and aligned chromosomes. In the TSD mice, the MII oocytes exhibited various morphologically abnormal spindles and misaligned chromosomes. Magnifications of the boxed regions are shown on the right of each main panel. Bar=20μm. **(B)** Percentages of oocytes with abnormal spindles and misaligned chromosomes in MII oocytes from the stationary, TSD and forced night activity groups. Data are presented as means±SEM of 3 independent experiments. Different letters denote statistically significant differences (P<0.05).

As shown in Figure 7 B, the GV oocytes from the TSD mice showed stronger red/green fluorescence ratio than the two control groups (0.75 versus 0.56 and 0.57, control; P<0.05), indicating an increased mitochondrial membrane potential (△ψm), corresponding to the compensatory increase in mtDNA number. In contrast, the relative red/green fluorescence ratio (△ψm) in MII oocytes from the TSD mice sharply decreased, compared to the two control groups (0.67 versus 1.03 and 0.91, control; P<0.05), indicating lower mitochondrial activity and a reduced △ψm. Collectively, these results indicate that maternal sleep deprivation causes a change of the mitochondrial membrane potential during oocyte maturation. The increased mtDNA number and the increased △ψm may represent compensation response of reduced quality of the GV oocytes, while the decreased mtDNA number and △ψm may represent an impaired oocyte quality.

### Maternal sleep deprivation is associated with abnormal spindles and misaligned chromosomes in oocytes

To further clarify the possible effects of maternal total sleep deprivation on oocyte quality and impaired embryonic development, ovulated MII oocytes were immune-labeled with α-tubulin antibody. As shown in Figure 8, most of MII oocytes in the stationary and the forced night activity control groups exhibited barrel-shaped spindles. However, a higher frequency of spindle defects was observed in ovulated MII oocytes from TSD mice (8.03±1.52%, n=75), compared with 3.9±0.97% and 4.37±0.98 % in the two control groups. The defective spindles include various malformed spindles, additional asters and disintegrated spindle poles (Fig. 8A). At the same time, the percentage of chromosome misalignment in TSD mouse oocytes also increased significantly (6.3±1.35 %, n=75), the lagging chromosomes and irregularly scattered chromosomes (Fig. 8 A-B), were shown to be considerably higher than those in the two control groups (2.2±0.67%,n=96 and 2.95±0.66%, n=83; respectively) (P<0.05) (Fig. 8 B).

## Materials and Methods

All chemicals and culture media were purchased from Sigma Chemical Company (St Louis, MO, USA) except for those specifically mentioned. All animal care and use procedures were conducted in accordance with the rules promulgated by the Ethics Committee of the Institute of Zoology, Chinese Academy of Sciences.

### Generation of mouse models

Six-week-old ICR mice acclimated in the animal facility for at least 1week prior to the experiments. Mice had free access to food and water. Throughout the experiment, the experimental room was maintained at an ambient temperature of 21℃, with lights on from 09:00 to 21:00. Mice were deprived of sleep via forced locomotion in an in-house-built sleep deprivation device, which consisted of a rotating drum (Supplement Figure 1). The drums were large, motorized, equipped with stainless-steel activity wheels with 22 cm outside diameter and 18 cm internal wheel width. The wheels were run through a computer-controlled motor mediated by a drive strip. The front and back panels of the drum are made of Plexiglas. A water bottle and a feeding RACH were mounted on the front panel. The lens of an infrared sensitive video camera was mounted 20 cm in front of the wheel. There was video surveillance throughout the deprivation.

Mice were randomly assigned to one of the following three groups. In the TSD (total sleep deprivation) group, the mice were submitted to 72 hours of total sleep deprivation, through slowly rotating the wheels at a constant speed (0.3m/min) [5,33]. In the stationary group, control mice were rested on the stationary wheels. The mice in the forced night activity group were placed in the drum that rotated at twice the speed (0.6m/min) during the night (12h per day), which correlates with the circadian activity phase. The mice in the forced night activity group were exercised at a higher intensity (total intensity same to the TSD group) and had more time to sleep (12h per day) compared to the TSD group. All mice were habituated to the drum for 2 days by placing them in the wheel, before starting the experiments.

### Oocyte collection and culture

The fully-grown GV-oocytes (>80 μm) were obtained by manual rupturing of antral ovarian follicles from ovaries of the stationary, TSD and forced night activity mice on day 3 (Supplement Figure 1). The GV oocytes were cultured in M2 medium supplemented with 200μm IBMX to prevent meiotic resumption. To collect the MII oocytes for following molecular, cellular, and biochemical analysis (Supplement Figure 1), the mice were removed from the drum at the end of the light period (prior to lights off, at 21:00) and superovulated by intraperitoneal injection of 10 IU PMSG (Tianjing Animal Hormone Factory); 48h later, the mice were injected intraperitoneally with 10 IU (HCG; Tianjing Animal Hormone Factory). The MII oocytes were recovered from the oviductal ampullae of the superovulated mice, 12-14h after HCG. And cumulus cells were removed by pipetting the M2 medium, containing 0.1% hyaluronidase.

### Fertility and natural ovulation analysis

Since the effect of sleep deprivation on mice is only a short time after the end of the sleep deprivation on day 3, we only used the females which were in first estrus phase after the end of the sleep deprivation on day 3 to determine the natural ovulation rate and the fertilization rate . In the mouse, the estrous cycle is divided into 4 stages (proestrus, estrus, metestrus, and diestrus) and repeats every 4 to 5 days unless interrupted by pregnancy, pseudopregnancy, or anestrus [34]. So we only detected the stage of estrous from day 3 to day 6 (Supplement Figure 1), using the vaginal cytology method [34]. That is to say, if the female mice were not in estrus phase after the day 6, we do not use them for mating and subsequent experiments.

Vaginal cycles in all females during and after the TSD period appeared normal and ranged in length from 4 to 5 days. Since the effect of sleep deprivation on mice is only a short time after the end of the experiment, we only used the females which were in first estrus phase after the end of the total sleep deprivation experiment to detect to detect the natural ovulation rate, the fertilization rate and the embryo development. To access the reproductive activity, seven individually housed stationary, TSD or forced movement mice were examined for the estrous cycle at the end of the sleep deprivation experiment from day 3 to day 6. Female mice in the estrus phase were mated with ICR male mice with known fertility. For the natural ovulation assay, females in estrus were crossed with ICR fertile male on day 3 to day 6 (>10 weeks of age). The next morning, female mice with plugs were killed, and zygotes were collected from the oviduct and counted. The zygotes were cultured in KSOM (K Simplex Optimization medium) at 37℃, 5% CO_2_ in air and examined for blastocysts after 5 days.

### Blood collection and hormone analysis

Mice removed from the drum at the end of the experiment were identified by the stage of estrous and taken for blood immediately. Vaginal cytology method was used to accurately identify all stages of the estrous cycle, as previously mentioned [34]. The mice were anesthetized by 2,2,2-tribromoenthanol (Sigma-Aldrich 75-80-9) injection, and then blood samples were taken via removing the eyeballs. All blood samples were clotted at 4℃ overnight, followed by low-speed centrifugation (876g for 20 min at 4℃) to separate the serum. Finally, the supernatant was stored at -80℃ for further analysis. Estrogen and FSH were measured in a commercial laboratory (Beijing Worth in statute of Biotechnology CO., Ltd.). The assay sensitivity was <0.5pg/ml (estrogen), 0.25IU/ml (FSH). And the intra- and inter-assay coefficients of variation were<10% and <15% (estrogen), 2.2-2.5% and 3.7-8.7% (FSH), respectively. Each serum sample was measured in triplicates.

### RNA sequencing and analysis

Eight GV-intact denuded oocytes were used for single-cell RNA sequencing per group. For each animal, the total time that elapsed between mouse euthanasia and snap-frozen samples did not exceed 30 min. RNA was amplified and sequenced by Annoroad Corporation (Beijing, China). RNA-seq data were deposited in the Genome Sequence Archive (Dataset CRA002676). The DNA library was constructed as described before [35]. The differently expressed genes (DEGs) were analyzed with edgeR package in R with default parameters. To eliminate the effects of gentle exercise on the transcriptome, we defined SETa as the 64 DEGs between the Total sleep deprivation (TSD) and stationary groups, SETb as the 109 DEGs between the stationary and forced movement groups; DEG_TSD_ defined the 42 DEGs obtained by subtracting SETb from SETa, namely the DEGs caused by total sleep deprivation (DEG_TSD_). Subsequent analysis was based on DEG_TSD_. The functional enrichment analyses of the significant DEG_TSD_, including GO, were conducted using DAVID (Version 6.8), with the cut-off criterion of adjusted p-value<0.05 and enrichment gene count>2. The GO biological processes and pathways were screened out. The figures were generated by ggplot2 package in R.

### Real-time quantitative PCR verification

RNA was extracted from 30 GV-intact denuded oocytes per group by using a RNeasy micro purification kit (Qiagen, Austin, TX, USA). Then, single-strand cDNA generated with the cDNA synthesis kit (Takara, Otsu, Japan) was performed, using poly T primers. These cDNAs were used as templates to amplify *Arntl2*, *Psg17*, *Tpm3*, *Zfp984*, *Atp5k, Usmg5, Prr18, Pr138, Ccnd2, Arid5b, Cd200* and *Mnd1*. The primers are shown in Table S1. *Gapdh* was chosen as the reference gene. The primers used for the amplification of *Gapdh* fragment are listed as follows, Forward: 5’-CCCCAATTGTGTCCGTCGTG-3’; Reverse: 5’- TGCCTGCTTCACCACCTTCT-3’. We utilized the Roche light Cycler 480 to perform the PCR, using SYBR Premix (Kangwei, Beijing, China). Each experiment was repeated at least 3 times. Relative gene expression was measured using real-time quantitative PCR and the 2(-Delta Delta C(T)) method.

### Determination of ROS products

The method for ROS detection was described previously [13,36]. Briefly, the GV and MII oocytes were incubated for 30 min at 37℃ in M2 medium (Sigma, USA) supplemented with 10mM carboxy-H2DCF diacetate (Cat#S0033, Beyotime). Then the oocytes were washed in M2 medium for at least 3 times, and mounted on glass slides. Fluorescence was measured using a Carl Zeiss LSM 780 confocal microscope with identical settings through the center of the oocyte at its largest nuclear diameter. The photographs were analyzed using ZEN (2012) software, measuring fluorescence intensity for each oocyte.

### Determination of mitochondrial DNA copy number by quantitative real-time PCR

The mitochondrial DNA (mtDNA) extraction and mtDNA quantitative real-time PCR procedures were described previously [36]. The procedure was briefly as follows. One oocyte was placed in a PCR tube containing 10μl of lysis buffer and incubated at 55℃ for 2h. Proteinase K was heat-inactivated at 95℃ for 10min. Then the samples were used for quantitative RT-PCR analysis. RT-PCR was performed using the ABI system and mouse mtDNA-specific primers: B6 forward, AACCTGGCACTGAGTCACCA, and B6 reverse, GGGTCTGAGTGTATATATCATGAAGAGAAT. To obtain standard curves, PCR products amplified with B6-for and B6-rev were ligated into T-vector. Seven 10-fold serial dilutions of purified plasmid standard DNA was used to generate a standard curve. The linear regression analysis of all standard curves for samples with copy numbers between 10 and 10^6^ showed a correlation coefficient greater than 0.98. All measurements were performed for three times.

### Immunofluorescence

For mitochondrion staining, oocytes were incubated in M2 medium containing 20 μm MitoTacker Red (Cat# M7512, Invitrogen, USA) for 20 min at 37℃. After washes, the oocytes were stained with Hoechst 33342 (10mg/ml) for 10 min at 37℃ and analyzed with a confocal laser scanning microscope (Zeiss LSM 780).

For mitochondrial membrane potential staining, the live oocytes were cultured in M2 medium containing 2μm JC-1 fluorochrome (Beyotime Institute of Biotechnology) for 30 min at 37℃, 5% CO_2_ in air [34]. After washes, oocytes were still incubated in M2 containing Hoechst 33342 (10ng/ml) for 10 min. Finally, the oocytes were analyzed by fluorescence microscopy (Zeiss LSM 780). The fluorescence intensity analysis of each oocyte was analyzed using the ZEN (2012) software.

For spindle and chromosome staining, oocytes were fixed in 4% paraformaldehyde for 30min, and blocked in 1% BSA-supplemented PBS for 2h at room temperature. Then, oocytes were stained overnight at 4℃ with 1:800 anti-α-tubulin-FITC antibody (Sigma, 76074). After three washes in washing buffer, chromosomes were stained with Hoechst 33342 for 20℃. Finally, oocytes were placed on glass slides with anti-fade mounting medium (DABCO) to delay photobleaching, and viewed under a confocal laser-scanning microscope (Zeiss LSM 780).

### Statistical analysis

At least 3 replications were performed for all experiments. Data are presented as mean±SEM, unless otherwise indicated. Statistical analysis was conducted by Student’s t test and ANOVA when appropriate with SPSS 27.0 software (SPSS, Inc.). P<0.05 was considered to be statistically significant.

## Discussion

Sleep insufficiency is an inescapable condition that is becoming increasingly common in modern society, and it causes several harmful consequences to health. Many studies have suggested that sleep deprivation causes reproduction ability decline for males [1], but there are a few studies related to its effects on female fertility. In the present study, we generated a sleep deprivation mouse model by depriving of sleep for 72h using the rotating drum method. We found that total sleep deprivation (TSD) affects E2 and FSH levels in mice. Although the TSD mice ovulated normally and eggs could be fertilized, but most fertilized eggs were stunted, resulting in fewer embryos being implanted. In addition, our data showed that TSD remarkably causes increased mitochondrial aggregation and dysfunction as well ROS production, and that TSD also evidently alters gene expression that is related to mitochondrial function.

### Maternal sleep deprivation disrupts sex hormone secretion

Female fertility depends on the coordinated development of oocytes and ovarian follicles, which are regulated by sex hormones, whereas sleep deprivation is closely related with sex hormones [38]. Sleep deprived adult female rats showed decreased estrogen concentrations compared to the control rats [7]. It has also been reported that sleep deprivation may regulate the release of ovarian hormones by altering hormonal neurochemical mechanisms [7]. Besides, sleeplessness in female shift workers suppresses melatonin production and causes hyperactivation of hypothalamic-pituitary adrenal (HPA), which leads to impaired secretion of sex hormones, anovulation and amenorrhea, failed embryo implantation. [19]. In our study, we found that compared to the stationary and the forced night activity control group the estrogen (E2) and FSH levels in TSD mice decreased significantly, especially at the proestrus of the estrus cycle. However, it is still unclear how sleep deprivation is related to sex hormone production. On one hand, it may be due to the inhibition of the hypothalamic-pituitary-adrenal (HPG) axis caused by elevated corticosteroids under certain stress conditions [24]. On the other hand, it was suggested that sleep deprivation could activate the serotonergic nervous system and promote the release of serotonin, therefore, serotonin suppressed FSH-induced estradiol release in large follicles [39]. Because serotonin inhibits estradiol production and decreases the expression of STAR protein in large follicles, estradiol concentrations may decrease during sleep deprivation [39].

As is known to all, gonadotropins, especially FSH, can enhance the production of estrogen in granulosa cells in pre-ovulatory follicles [40]. Besides, decreased levels and irregular cyclical patterns of FSH and estradiol are known to have harmful effects on human sexual function and fertility [41]. We observed that TSD caused decreased fertility. Therefore, we propose that sleep deprivation in females is related to low E2 and FSH levels, which may compromise follicle function and oocyte quality, leading to female subfertility.

### Maternal sleep deprivation alters expression of genes, especially those related with mitochondrial functions

Since ovulation was not affected but embryo implantation and fertility were reduced in TSD mice, it was highly possible that oocyte quality was impaired. To reveal the mechanisms, the gene expression profiles of oocyte and their regulatory mechanisms in TSD model mice were studied by RNA-Seq technique for the first time. A total of 42 DEGs were identified, which were verified by random sampling RT-PCR. Most of the DEGS identified were related with mitochondria, such as *ATP5k*, *UMSg5* and so on. ATP5k, a protein required for maintaining the ATP synthase dimeric from [26], is involved in the biological process of “oxidoreductase activity”, and its down-regulation suggests the energy supply was decreased in the oocytes of TSD mice. Umsg5 plays roles in Complex V (ATP synthase), whose knock-out reduces ATP synthesis and mitochondria dysfunction [42]. In our study, we demonstrated a significant decrease in *ATP5k* and *Umsg5* gene expressions in sleep-deprivation mice. This decrease may reflect reduction of energy supply and mitochondria dysfunction caused by total sleep deprivation in mouse oocytes. Apart from the mitochondria associated the DEGs above, several other genes were identified, such as *TPM3*, *Cyclin D2*, and *Rp138*, which were previously shown to be associated with the regulation of oocyte meiotic maturation [27,28,43]. The underlying mechanisms of most of the DEGs associated with TSD are still unknown, and further studies are required to understand this phenomenon.

### Increased ROS in oocytes of TSD mice

Reactive Oxygen Species (ROS) is by-product of biological aerobic metabolism, and is a general term for a class of oxygen-containing and active substances. [18]. ROS at the physiological level play a key role in the signaling pathway mediating meiotic resumption and cell proliferation, while excessive and prolonged levels of ROS can cause oxidative stress which is highly damaging [44]. Oxidative stress is the imbalance between the production of reactive oxygen species in the body and the antioxidant system, resulting in the irreversible damage of biological macromolecules (lipids, proteins and DNA) in the body [45]. Sleep deprivation could impair the transport of electrons from complex I to complex III, resulting in an over production of ROS and leading to oxidative damage [16]. It has been assumed that ROS and oxidative stress in liver, spleen, nose, ganglion, heart, and plasma were the cause of the sleep deprivation syndrome [18]. Moreover, repeated extended durations of sleep loss increase superoxide production and acetylation of mitochondrial protein in Locus ceruleus neurons (LCn), and thus apoptosis is activated and LCns are lost [3]. In addition, melatonin is the major synchronizing agent of the circadian sleep pattern, and it also cleans up free radicals by regulating expression of antioxidants and reducing pro-apoptotic proteins, such as Bcl-2-associated X protein (BAX) [46]. Melatonin plays an important role in mouse and human oocyte maturation, fertilization and early development of embryonic tissue as well [47,48]. SD reduced the secretion of endogenous melatonin, which limits the levels of follicular melatonin and thus exposing the oocyte to high levels of ROS, which may reduce oocyte quality in infertile women [49,50]. IVF patients with sleep disorders may benefit from melatonin supplementation by improving the quality, maturation and number of oocytes retrieved and thus improving the quality and number of embryos [46]. The present study showed significantly increased levels of ROS in the GV and MII oocytes from the TSD mice compared to those in the stationary control and the forced movement control groups, which is in agreement with the reports above in other systems. High metabolism is required to maintain electric potentials during wakefulness, which requires a large amount of oxygen, resulting in production of ROS in large quantities [18]. Once mitochondrial and cytoplasmic antioxidant systems are overwhelmed by ROS, oxidative damage occurs [16]. The oocytes and embryos of mammalians are extremely sensitive to oxidative stress, which can destroy oocyte’s function by oxidating RNA, DNA, proteins; damaging the embryo development; and promoting embryo fragmentation [50]. Thus, sleep deprivation increases ROS production during mouse oocyte meiotic maturation, which might disrupt the quality of oocyte and early embryo development in mice.

### Maternal sleep deprivation disrupts mitochondrial function in oocytes

In humans and animals, sleep deprivation could lead to mitochondrial dysfunction and oxidative stress [15]. It was demonstrated that sleep deprivation was able to cause mitochondrial dysfunction, as marked by decreased activity of complex I-III, II and II-III of the mitochondrial electron transport chain, effects which could in turn be involved in increased oxidative damage [16]. Mitochondria are the most numerous organelles in mammalian oocytes and early embryos, and dysfunction or abnormality of mitochondria in oocytes may be a key determinant of embryo development [10,13]. Factors such as mitochondrial DNA copy numbers, <u>mitochondrial transmembrane potent</u>ial (ΔΨm), stage-specific spatial distribution and structural dysfunctions may affect the developmental competence of oocytes and preimplantation embryos [10]. To evaluate the effects of sleep deprivation on mitochondrial function, we evaluated the mtDNA copy number, mitochondrial distribution, and ΔΨm in oocytes.

It has been demonstrated that low mtDNA numbers are related to arrest of preovulatory meiotic maturation, fertilization failure, and even ovarian dysfunction [29]. It was demonstrated that mtDNA copy number is increased in oocytes from older women or DEM women [11,29]. Our results showed an increase in the number of copies of mtDNA in the GV oocytes from TSD mice compared to that in the stationary and forced movement control oocytes (Fig.6A). Unusually high mtDNA copy numbers may be associated with high ATP supply. Such an increase in mtDNA copy number of the TSD mice GV oocytes may be ascribed to a compensatory phenomenon to ensure the production of sufficient ATP in the event of increased demand or respiratory chain dysfunction caused by TSD. When the supply of ATP exceeds cellular demand, the levels of oxidative free radical production rises, which can lead to cell death of zygotes, through disrupting activity and structures of mitochondrial [51]. Strikingly, our results showed that the mt DNA copy number of TSD mice MII oocytes was significantly lower than that in the stationary and forced movement control oocytes (Fig.6 B). The loss of mtDNA at the MII stage may be caused by OS impairment and increase in mitochondrial degradation or autophagy [13,51]. Studies have shown that acute sleep deprivation significantly reduces levels of important mitochondria in the hippocampus [52]. In addition, low mtDNA copy numbers in human MII oocytes, may harm the developmental ability of oocytes and the preimplantation-stage embryos, since low mt DNA copy numbers could adversely affect the metabolic capacity [10]. Next, we analyzed the stage-specific distribution of mitochondria in mouse oocytes. TSD contributes to an inadequate translocation of active mitochondria during mouse oocyte meiotic maturation and mitochondrial clustering was clearly increased in TSD mice oocytes compared with the oocytes from the stationary and the forced movement control groups (Fig.7). During oocyte maturation, the distribution of mitochondria goes through stage-specific changes. It is commonly believed that stage-specific spatial remodeling of mitochondria may allow higher levels of environmental ATP to occur in cytoplasmic regions where stage-specific activities such as GV breakdown and spindle formation takes place which have higher energy demands [10]. Thereby, inadequate mitochondrial redistribution and heterogeneous mitochondrial fragmentations may result in insufficient energy provided by the organelles, which may be a cause of developmental retardation in TSD mice.

In the oocyte mitochondria, energy is stored as mitochondrial membrane potential (ΔΨm), the energy stored in which drives the conversion of ADP to ATP by the operation of the respiratory chain enzymes [53]. Higher polarized mitochondria were reported to localize around the periphery cytoplasm in the MII oocytes, which presents high mitochondrial membrane potential and high ATP supply [54]. JC-1 is widely used as a ΔΨm-specific probe, which can form aggregates (red fluorescence) from monomers (green fluorescence). The red fluorescence represents the higher ΔΨm, and the green fluorescence represents lower ΔΨm. The relative ratio of red to green fluorescence intensity is often utilized to indicate the functional status of mitochondria [31]. Interestingly, our present study shows that the GV oocytes from TSD mice transiently showed higher mitochondrial activity, which may be a compensatory phenomenon to ensure adequate energy supplement, as sleep deprivation induced elevated energy expenditure (Fig. 8). Notably, increased ΔΨm was also found in the GV oocytes from the insulin resistance mice, which was associated with poor oocyte quality [13]. MII oocytes from TSD mice showed lower ΔΨm (Fig. 8). Acton et, al. reported that fertilization failure and embryo arrest was related to a reduction in the ratio of high-to-low polarized mitochondria in MII oocytes [32]. These observations indicate that TSD can cause injuries to oocyte mitochondrial membranes, thereby impairing the activity of mitochondria.

Taken together, maternal sleep deprivation leads to the occurrence of oxidative damage, promotes mitochondrial dysfunction and generation of more reactive oxygen species, resulting in a vicious cycle and leading to poor oocyte quality and impaired developmental potential of embryos.

### Maternal TSD compromises nuclear dynamic events in oocytes

Oocyte mitochondrial defects, such as altered ΔΨm, altered cristae structure, altered mitochondria localization and reduced ATP production, will disrupt mitotic and meiotic spindles, inhibiting oocyte maturation and early embryo development [56]. Our cytological analysis revealed that maternal total sleep deprivation leads to an increased frequency of abnormal meiotic spindle formation and chromosome abnormalities in mouse oocytes (Fig. 9). First, increased ROS and increased oocyte susceptibility to ROS could cause spindle instability, chromosomal misalignment and was thought to be a cause of reduced developmental competence of oocyte [57]. Second, it is revealed that deterioration of mitochondria in oocytes decreases ATP production and disrupts the formation of meiotic spindle [36]. In addition, the interaction of mitochondria with spindle poles was found to reduce spindle rotation and promote spindle alignment in the fission yeast *Schizosaccharomyces pombe* [58]. And mitochondria distribution affects spindle formation [10]. In conclusion, mitochondrial dysfunction plays a crucial role in meiotic defects in TSD oocytes, and these results have important implications for understanding the harmful embryo development in TSD mice.

In conclusion, our study indicates that total sleep deprivation is detrimental to female fertility in mice by affecting sex hormone secretion and altering the oocyte transcriptome. It also leads to OS in the GV and MII oocytes and impairs mitochondrial function, which affects oocyte quality and embryo development.

## ACKNOWLEDGMENTS

This study was supported by The National Natural Science Foundation of China [82001541]; the National Natural Science Foundation of China [31530049]; the Basic Research Key Project of Shenzhen Science and Technology Innovation Commission [JCYJ20200109140623124].

## AUTHOR CONTRIBUTIONS

Q.-Y. S., W.-P. Q. designed and conceived the experiments; Z.-Y. Y., Q.-X.L. and Q.-Z. performed the experiments; Z.-Y. Y. wrote and all the authors reviewed the manuscript. C.-H. Z., and J. Q. provided professional advice on experimental design and paper writing. H. S. and Q.-Y. S edited the manuscript. Y.-L., Q.-R. M., J.-L., Y.-H. L., Y.-P. C., provided assistance with experiment performance.

## DATA AVAILABILITY STATEMENT

The data used to support the findings of this study are included within the article. All data generated or analyzed during this study are included in this article.

## DECLARATIONS

### ETHICS APPROVAL AND CONSENT TO PARTICIPATE

Not applicable.

### CONFLICT OF INTERESTS

The authors declare that there are no conflict of interests.

### CONSENT FOR PUBLICATION

Not applicable.

---

**Supplement Figure 1.**
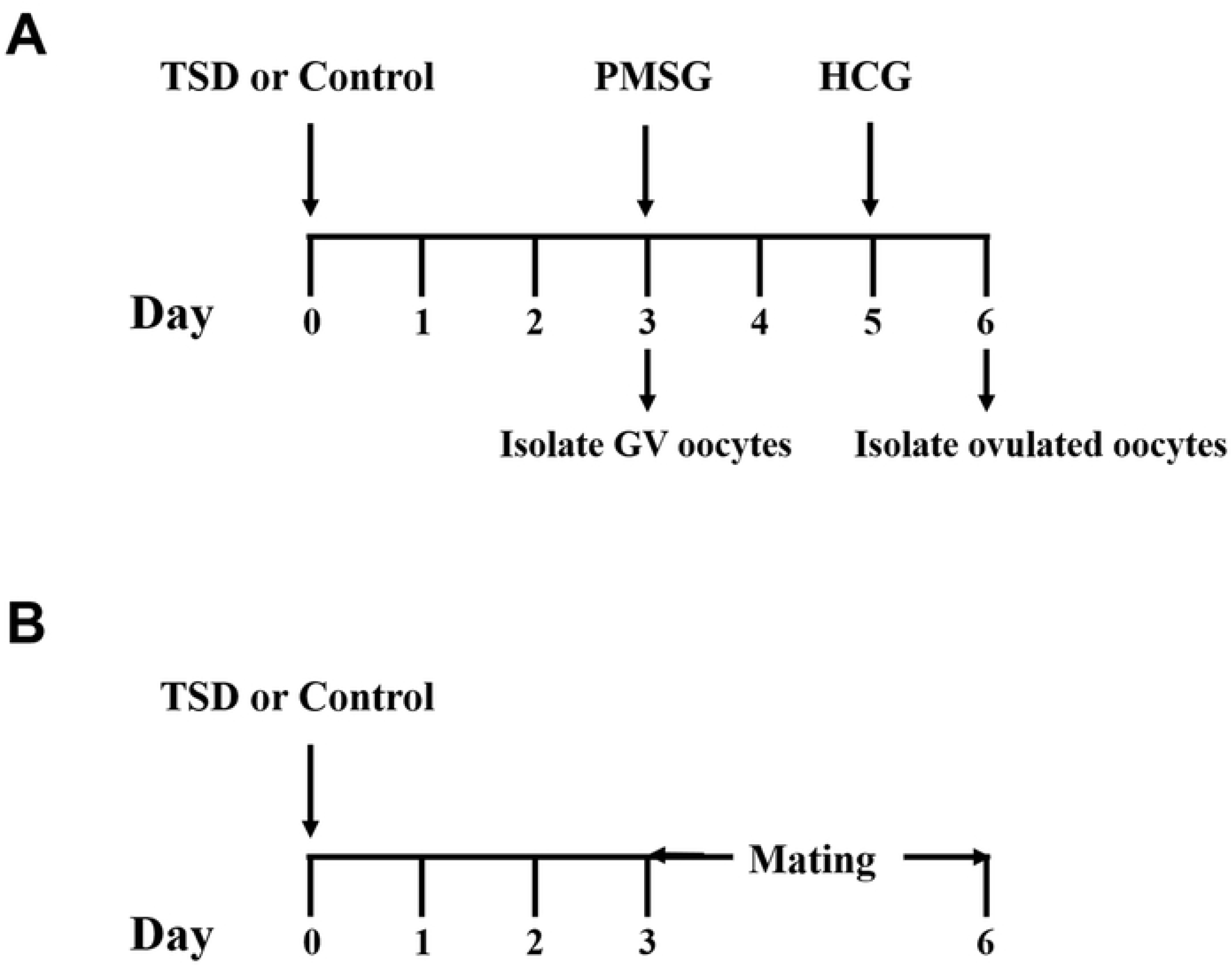
Schematic illustration of generation of mouse models and oocyte collection. (A) Female mice were deprived of sleep via forced locomotion on day 0. There days later, part of the mice were identified by the stage of estrous and taken for blood immediately. Part of the mice were sacrificed to collect the GV oocytes. To retrieve ovulated oocytes for following molecular, cellular and biochemical analysis, part of the mice were administrated with 10 IU PMSG (d 3). These mice were injected with 10 IU HCG 2 d after PMSG. At 13.5 h after HCG administration, oocytes were collected from oviductal ampullae (d 6).

## Notes

### Competing Interest Statement

The authors have declared no competing interest.

